# Alternative splicing tunes sodium channels to support channel- and neuron-specific effects

**DOI:** 10.1101/2020.09.30.320788

**Authors:** Gabriele Lignani, Andrianos Liavas, Dimitri M Kullmann, Stephanie Schorge

## Abstract

Neuronal excitability is tightly regulated, requiring rapidly activating and inactivating voltage-gated sodium channels to allow accurate temporal encoding of information. Alternative splicing greatly broadens the repertoire of channels, but the adaptive significance of this phenomenon is incompletely understood. An alternative splicing event that is conserved across vertebrates affects part of the first domain of sodium channels and modulates their availability after inactivation. Here we use this conserved splicing event to ask whether this modulation has consistent effects in different neuronal backgrounds, or whether a conserved splicing event can be exploited to produce distinct effects in different cell types. We show that the consequences of alternate splicing of human Nav1.1 and Nav1.2 for neuronal activity depend on whether they are expressed in the cell types where they normally predominate (interneurons or excitatory neurons, respectively). Splicing in the ‘adult’ isoform in both channels is sufficient to slow action potential rise times in all neurons. However, changes to both action potential half width and maximal firing rate are specific to cell type and channel, with each channel appearing tuned to mediate effects in its predominant neuronal background. Finally, we use dynamic clamp to demonstrate that alternative splicing in Nav1.1 changes how interneurons fire during epileptiform events. Our data show that, for sodium channels, despite conserved amino acid changes and similar effects on channel gating, alternative splicing has distinct impacts on neuronal properties, thus highlighting how closely sodium channels are tuned to distinct cellular backgrounds.

## Introduction

Mutations in sodium channel genes are increasingly associated with different neurological disorders, and there is growing consensus that mutations which lead to gain of function (GOF) or loss of function (LOF) can lead to clinically distinct disorders, and this dependent both on the type of gene and the channel targeted. For example SCN2A, which is thought to dominate early in development at axon initial segments (AIS) in excitatory neurons, and plays an important role in dendritic excitability later in life (Spratt et al., 2019), is associated with severe seizure disorders when mutations increase function, but LOF mutations in the same gene are one of the strongest risk factors for autism (Ben-Shalom et al., 2017)(https://gene.sfari.org/). In contrast a growing genotype-phenotype spectrum suggests that LOF mutations in SCN1A are closely linked to the severe epilepsy Dravet Syndrome, while GOF in this channel is more associated with migraine (Brunklaus et al., 2020a). The link between LOF and increased seizures, is thought to be because loss of this channel may predominantly affect inhibitory interneurons (Yu et al., 2006).

The increasing availability of genetic data linked to high quality clinical findings, means that a growing number of mutations producing similar functional consequences in different sodium channels have been reported. These mutations can lead to different disease manifestations, but these manifestations are consistent with the mutations having similar impacts on the channels themselves, i.e. a LOF mutation linked to Dravet syndrome in SCN1A is likely to also lead to a LOF impact in SCN2A, but LOF in SCN2A would be expected to be associated with risk of autism rather than epilepsy, as seizures are more associated with GOF in SCN2A (Brunklaus et al., 2020b).

Highly conserved splicing in the first domain of sodium channels imposes functionally conserved effects on channel behavior in neuronal sodium channels (Liavas et al., 2017). The alternate splice variants are conventionally designated ‘Adult’ or ‘A’, and ‘Neonatal’ or ‘N’, affecting the 5^th^ exon of *SCN1A* (Nav1.1 6A or Nav1.1 6N) or the homologous 6^th^ exon of *SCN2A* (Nav1.2 6A or Nav1.2 6B). By comparing the consequences of expressing two splice variants, it is possible to ask whether a homologous change, with conserved functional consequences in different channels, has evolved to produce conserved changes in neuronal activity, or whether the diversity of channels and neurons allows even a conserved change to be exploited for different consequences in difference cell types.

We show that the underlying change in channel availability caused by splicing is sufficient to change action potential rise time in trains of rapid stimuli, and this is seen using either channel and in both excitatory and inhibitory neurons. However, additional effects of splicing are determined both by the channel type used, and on the neuronal background in which the channel is expressed. These data not only reveal a consistent impact of splicing on action potential kinetics, but also demonstrate how exquisitely tuned sodium channels are to different neuronal properties.

## Materials and methods

### DNA constructs and Cloning

Human Nav1.1 cDNAs in the pcDM8 vector were transformed into TOP10/P3 cells, and Nav1.2 cDNAs were in pcDNA3 vectors and transformed into Stbl3 cells, as described previously (Liavas et al., 2017). Site-directed mutagenesis was performed in Nav1.1 to introduce a single amino acid change (F383S) in order to confer TTX resistance according to previous studies (Bechi et al., 2012; Cestèle et al., 2013) using the QuikChange II XL kit according to the manufacturer’s instructions (Stratagene, CA). The homologous mutation was also performed in Nav1.2 (F385S)(Rush et al., 2005). Successful mutagenesis was confirmed by DNA sequencing.

### Hippocampal neuron culture and transfection

Hippocampal neurons were isolated from P0 GAD67-GFP knock-in mouse pups (Tamamaki et al., 2003), where interneurons can be visually distinguished by GFP expression as previously described (Kaech and Banker, 2006). The pcDM8-hNav1.1 5A/5N or pcDNA3-hNav1.2 6A/6N DNA vector was co-transfected with a reporter Red Fluorescent Protein (RFP)-carrying plasmid under a beta-actin promoter in a 5 : 1 molar ratio. The neurons were transfected on day 4 after plating by magnetofection with NeuroMag according to the manufacturer’s instructions (OZ biosciences). Recordings were performed 3-6 days after transfection. For recordings from interneurons, cells patched showed co-localization of both green (indicating interneurons) and red (indicating successful transfection with the sodium channel plasmid) fluorescence.

### Whole cell patch clamp recordings in neurons

For current-clamp recordings of transfected neurons, the internal solution contained (in mM): 126 K-gluconate, 4 NaCl, 1 MgSO_4_, 0.02 CaCl_2_, 0.1 BAPTA, 15 Glucose, 5 HEPES, 3 ATP-Na_2_, 0.1 GTP-Na, pH 7.3. The extracellular (bath) solution contained (in mM): 2 CaCl_2_, 140 NaCl, 1 MgCl_2_, 10 HEPES, 4 KCl, 10 glucose, pH 7.3. D-(-)-2-amino-5-phosphonopentanoic acid (D-AP5; 50 μM), 6-cyano-7-nitroquinoxaline-2,3-dione (CNQX; 10 μM) and picrotoxin (PTX; 30 μM) were added to block synaptic transmission. Tetrodotoxin (TTX; 1 μM) was added to block endogenous sodium channels, allowing isolation of transfected TTX-resistant sodium currents. Experiments were performed at room temperature (22-24°C). Neurons with unstable resting potential and/or bridge-balance >15 MΩ were discarded. Bridge balance compensation was applied and the resting membrane potential was held at −70 mV. Action potentials were evoked by injecting 10 ms long depolarizing current steps of increasing amplitude. For the firing frequency reliability protocol, a 110% value of the threshold current was delivered in 11 consecutive pulses with increasing frequency in each series (33 – 90 Hz). All recordings and analysis for neurons were carried by a researcher blinded to the isoform expressed. Recordings were acquired using a Multiclamp 700B amplifier (Axon Instruments, Molecular Devices, Sunnyvale, CA, USA) using in house software written in LabVIEW 8.0 (DMK), filtered at 10 kHz and digitized at 50 kHz.

### Analysis of single action potential parameters

The single action potential shape parameters were derived using pClamp (Molecular Devices) and analyzed with Prism (GraphPad Software, Inc.). A phase-plane plot of the first action potential elicited after sufficient depolarization with current steps was obtained for each cell by plotting the time derivative of voltage (dV/dt) versus the voltage. This allowed identification of the AP voltage threshold, peak and amplitude as well as the maximum rising and depolarizing slopes (Bean, 2007). The action potential threshold was defined as the voltage at which dV/dt exceeded 10 mV/ms, similar to other studies (Pozzi et al., 2013).

### Dynamic clamp recordings in neurons

Recordings were carried out as described previously (Morris et al., 2017). Briefly, current traces in voltage-clamp configuration in the presence of 4AP were recorded holding neurons at −70mV, the resulting current traces were converted in conductance (G=I/V). Using Signal dynamic clamp software in conjunction with CED Power 1401-3 (CED, Cambridge Electronic Design Limited) the conductance traces were used to inject currents in neurons (in 1μM TTX), in the current clamp configuration. During recordings, the voltage of the patched neurons was read in real time, and used to calculate the current to be injected from the 4AP conductance trace. In order to compare different cells, the conductance threshold was calculated in each neuron prior to each dynamic clamp experiment using AMPA conductance steps (E_rev_=0mV; τ=1ms; ΔG=1nS), and the injected epileptiform conductance traces were scaled to this threshold. To scale each neuron a 15% of the conductance threshold was injected inn both inhibitory and excitatory neurons. This value has been used because neurons reached the plateau and their maximal firing rate at this percentage (Figure 1).

**Figure 1:**
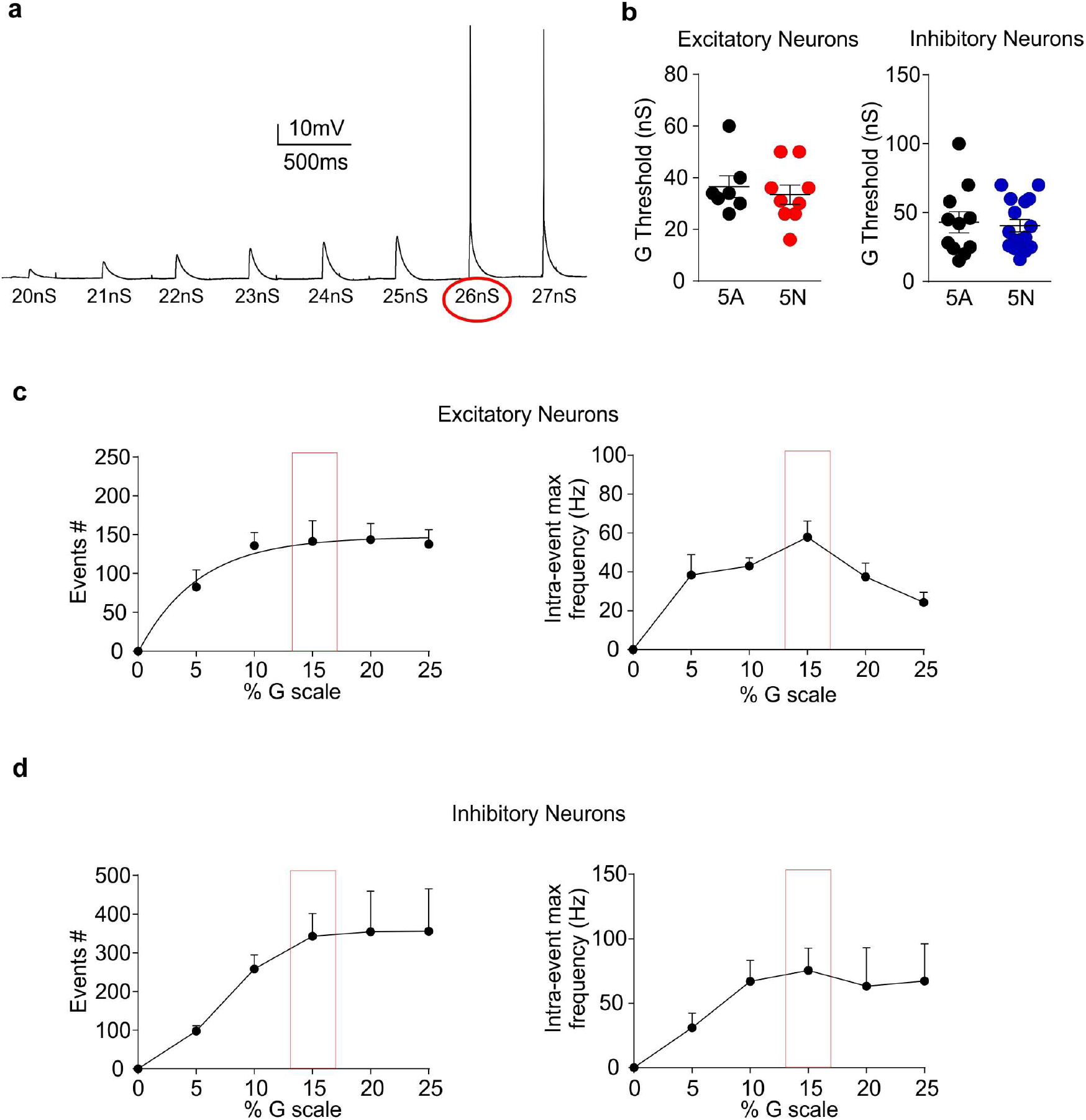
Dynamic clamp experiments to set the percent of conductance threshold for activity clamp. **(a)** Representative protocol of dynamic clamp AMPA conductance steps (E_rev_=0mV; τ=1ms; ΔG=1nS). The red circle is the conductance threshold in this specific experiment (repeated 10 times for each neuron recorded). **(b)** Conductance threshold is similar in neurons expressing either 5N or 5A splicing isoform of Nav1.1 in both inhibitory and excitatory neurons. **(c and d)** Number of events and intra-event maximal frequency in excitatory and in inhibitory neurons, respectively, against the percent of conductance threshold. The red squares represent the percent of conductance threshold used for all the dynamic clamp experiments.

### Statistical analysis

Results are shown as mean ± s.e.m. Data were tested for normality using the Shapiro-Wilk test. Normally distributed two sample groups were compared by Student’s unpaired two-tailed t-test, at a significance level of P < 0.05. Sample groups without a normal distribution were compared using Fisher’s exact test (two-tailed). For comparisons of more than two sample groups, ANOVA was used, followed by either the Bonferroni method or Dunnett’s test. Statistical analysis was carried out using Prism, Origin (OriginLab) or SPSS (IBM).

## Results

### Functional comparison of sodium channel variants in neurons

In HEK cells, alternate splicing of the same region of Nav1.1, NaV1.2 or Nav1.7 (encoded by *SCN1A, SCN2A* and *SCN9A* respectively) has a conserved effect on channel availability during trains of depolarizing steps (Liavas et al., 2017). Extrapolating these findings to neurons presents two challenges. First, multiple voltage-gated sodium channels are expressed, and second, the inhomogeneous distribution of channels in different compartments, including the axon and dendrites, precludes good voltage control.

To overcome these challenges, we characterized the effects of the splice variants in channels engineered to have reduced sensitivity to TTX block (see methods), and recorded from neurons transfected with individual variants in the presence of 1 μM TTX, thereby allowing the contributions of the transfected variants to be isolated from endogenous sodium channels. Furthermore, we used proxy measures of sodium channel activity and availability in the current clamp configuration. While current clamp cannot give a direct measure of the current density produced by the different channel variants, the density of sodium channels is correlated to the rising slope of action potentials, which is increased when more channels are available, and slowed when fewer sodium channels are available due to accumulation in inactivated states during trains of rapid firing (Carter and Bean, 2011). Changes to the falling phase of action potentials are largely set by the presence of different potassium channels in different neurons, and consequently of less utility in assessing sodium channel behavior (Bean, 2007).

### Effects of splicing in Nav1.2 in excitatory neurons

Nav1.2 is thought to be predominately expressed in excitatory neurons, particularly in early development, and its location and function change in adulthood (Oliva et al., 2012; Spratt et al., 2019). To isolate excitatory neurons, we transfected N or A splice variants of Nav1.2 into primary hippocampal cultures prepared from GAD67-GFP knock-in mice (Tamamaki et al., 2003) which allows excitatory neurons to be discriminated from inhibitory neurons. In the presence of TTX, excitatory neurons expressing TTX-resistant splice variants of Nav1.2 had similar AP thresholds, rising slopes, and amplitudes, as well as passive membrane properties (RMP, input resistance), suggesting a similar density of channel expression for both variants (Table 1). We applied depolarizing pulses at different frequencies and examined both the ability to generate APs (Figure 2A-C) and the AP shape (Figure 2D).

**Table 1.**
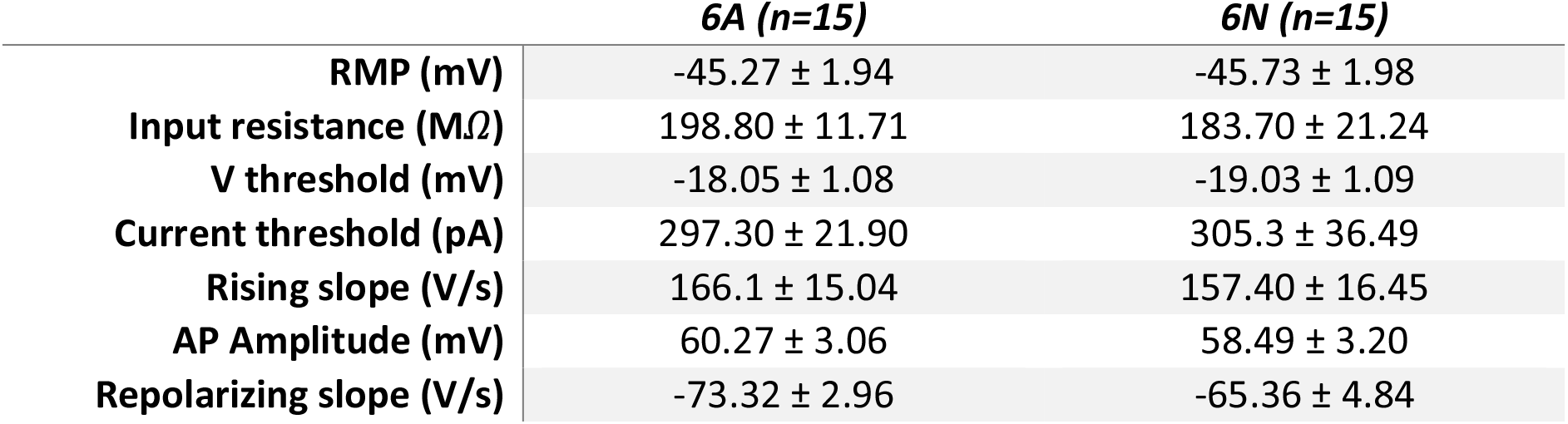
Membrane properties of excitatory neurons transfected with splice variants of NaV1.2 (mean ± s.e.m.)

|  | **6A (n=15)** | **6N (n=15)** |
| --- | --- | --- |
| **RMP (mV)** | -45.27 ± 1.94 | -45.73 ± 1.98 |
| **Input resistance (MΩ)** | 198.80 ± 11.71 | 183.70 ± 21.24 |
| **V threshold (mV)** | -18.05 ± 1.08 | -19.03 ± 1.09 |
| **Current threshold (pA)** | 297.30 ± 21.90 | 305.3 ± 36.49 |
| **Rising slope (V/s)** | 166.1 ± 15.04 | 157.40 ± 16.45 |
| **AP Amplitude (mV)** | 60.27 ± 3.06 | 58.49 ± 3.20 |
| **Repolarizing slope (V/s)** | -73.32 ± 2.96 | -65.36 ± 4.84 |

**Figure 2:**
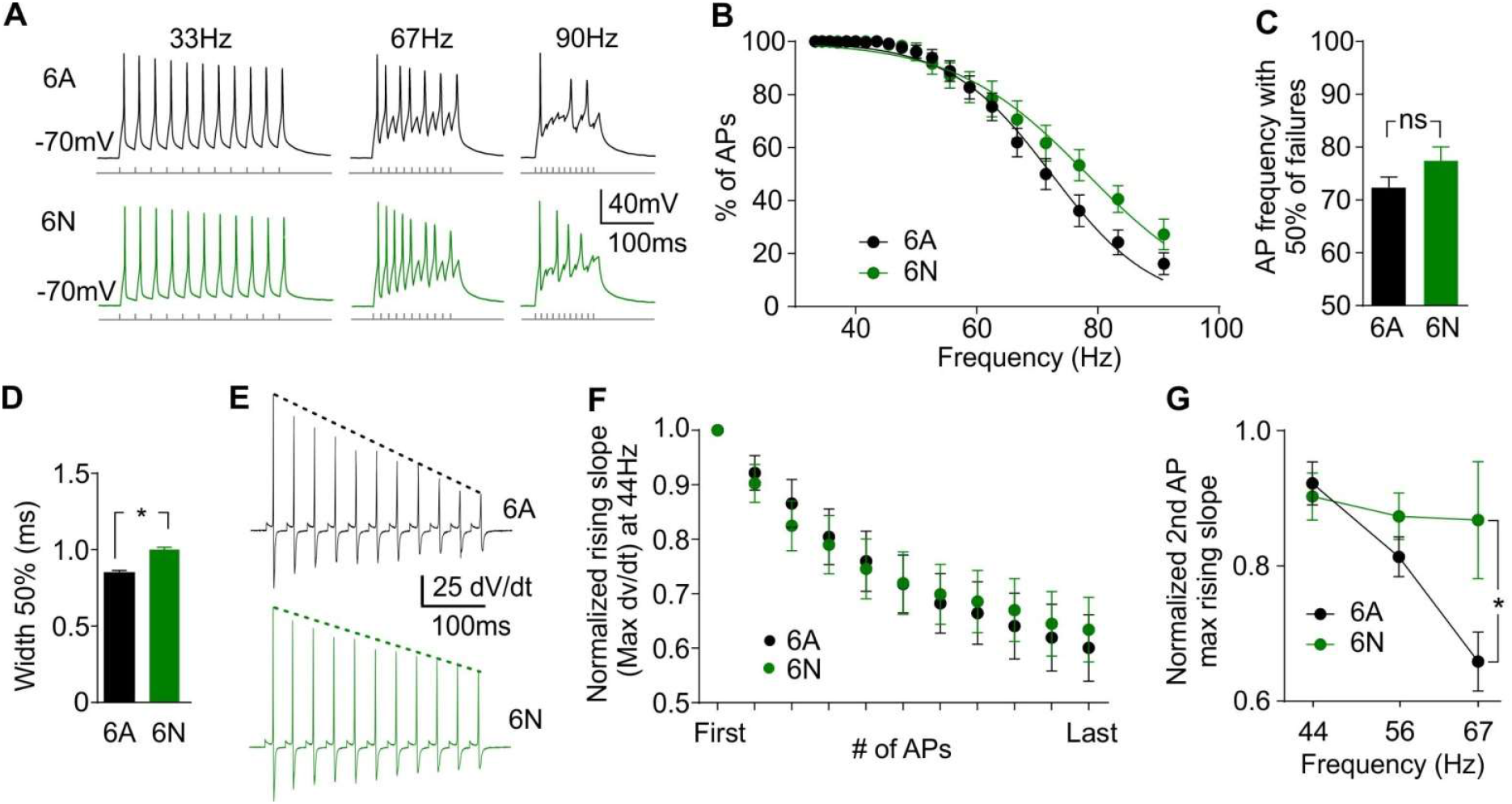
Splice variants of Nav1.2 in excitatory neurons have conserved effects on rising phase of action potentials during fast trains but not on maximal firing rates. **(A)** Representative traces from excitatory neurons transfected with A (top, black) and N (bottom, green) splice variants of Nav1.2. **(B)** Excitatory neurons expressing the N variant of Nav1.2 are able to sustain slightly, but not significantly, higher rates of firing than those expressing the A variant. **(C)** The rate at which excitatory neurons fired action potentials (APs) for 50% of stimuli, as calculated by average of exponential fits to individual cells, was not different (ns P > 0.05, unpaired two-tailed Student’s T test). **(D)** Expression of the A variant of Nav1.2 reduces the half width of APs in excitatory neurons. The mean half width was shorter for neurons expressing the A variant compared to the N variant (* P = 0.011; unpaired two-tailed Student’s T test). **(E)** The second differential of the APs evoked by representative series of steps from an excitatory neuron expressing the A (top, black), or N (bottom, green) variant of Nav1.2, showing marked drop in the speed of the rising slope for both variants. Traces are from stimuli at 44 Hz. **(F)** During trains of stimuli neither isoform is able to maintain rapid rising phases of APs in excitatory neurons. **(G)** At higher rates of firing the reduction in rising slope for A variant is evident after a single AP. Despite the smaller intrinsic differences in splice variants of Nav1.2 compared to Nav1.1, the Nav1.2 variants confirm that at high frequencies, the N-containing channels are more able to support fast rising phases. Data show the change in rising slope as measured from the second derivative for those excitatory neurons that fired in response to both the first and second stimuli. Cells which failed to fire APs were excluded (A n = 10; N n = 9). At 67 Hz the difference was significant (* P < 0.05; two-way ANOVA followed by Bonferroni’s multiple comparisons test).

A previous study showed that animals engineered to express only the A variant of SCN2A throughout development exhibited multiple changes in immature neurons, including an increase in firing rate and a shortened AP half width, but these differences were obscured as neurons matured (Gazina et al., 2015). Similar to the *in vivo* effects, we found in our system that the A variant of Nav1.2 also produced APs with shorter half widths (Figure 2D).

In HEK cells the most consistent functional consequence of alternative splicing was reduced availability of the A variants to activate during trains of stimuli (Liavas et al., 2017). In three different channels, incorporation of the A exon led to reduced channel availability after short depolarizing pulses. In neurons in the current clamp configuration, reduced channel availability would be consistent with splicing changing the ability of neurons to fire APs in response to rapid stimuli. For example, by reducing channel availably after short depolarizations, inclusion of the A exon might reduce the maximal firing rate, or progressively slow the rising phase of APs which can be limited by number of available channels. To test whether splicing was sufficient to change the availability of sodium channels during trains of rapid stimulations, we compared excitatory neurons expressing both splice variants of SCN2A, and asked whether AP parameters were affected during trains of stimulations.

Although there was a trend for the N variant of Nav1.2 to support firing at higher rates than the A variant, the difference did not reach significance (Figure 2A-C). This suggests that the alteration in availability is not sufficient in these neurons to disrupt initiation of regenerative APs. Both neonatal and adult variants of Nav1.2 showed pronounced slowing of AP rise-times during trains, consistent with reduced channel availability (at a frequency of 44 Hz; Figure 2E, F). The only robust difference between splice variants when examining AP rise times was observed with short inter-stimulus intervals (Figure 2G). The slower rise time for the A variant at 67 Hz is qualitatively consistent with data from HEK expression which also demonstrated more pronounced effects of splicing after shorter intervals (Liavas et al., 2017).

### Splicing in Nav1.1 changes channel availability and spike reliability in interneurons

Although splicing in all three channels studied in HEK cells imposed conserved changes on channel availability, the background rates of recovery were specific for each channel, suggesting these channels may be tuned to the neuronal types in which they are thought to predominate. Specifically, Nav1.1 and Nav1.2 channels underlie excitability of interneurons and principal neurons respectively (Ogiwara et al., 2007; Yu et al., 2006). We therefore asked how splice variants of Nav1.1 affect the firing of interneurons. As with Nav1.2, in the presence of TTX, neurons expressing TTX-resistant splice variants of Nav1.1 had similar passive membrane properties (Table 2).

**Table 2.** Membrane properties of inhibitory neurons transfected with splice variants of NaV1.1 (mean ± s.e.m.)

|  | **5A (n=15)** | **5N (n=13)** |
| --- | --- | --- |
| **RMP (mV)** | -57.80 ± 1.84 | -56.85 ± 1.76 |
| **Input resistance (MΩ)** | 146.2 ± 27.80 | 131.00 ± 15.64 |
| **V threshold (mV)** | -25.49 ± 1.17 | -24.37 ± 1.28 |
| **Current threshold (pA)** | 380.00 ± 66.63 | 312.30 ± 54.94 |
| **Rising slope (V/s)** | 82.82 ± 9.32 | 94.06 ± 9.99 |
| **AP Amplitude (mV)** | 36.96 ± 3.01 | 39.45 ± 3.24 |
| **Repolarizing slope (V/s)** | -47.36 ± 2.95 | -51.19 ± 2.31 |

We applied a similar series of depolarizing pulses at different frequencies in the presence of TTX (Fig. 2A-C). In striking contrast to Nav1.2 in excitatory neurons, splicing in Nav1.1 had a major effect on action potential reliability in interneurons, especially pronounced at intermediate frequencies (Figure 3B, C). It however had no significant effect on the half-width of APs in inhibitory neurons (Figure 3D).

**Figure 3:**
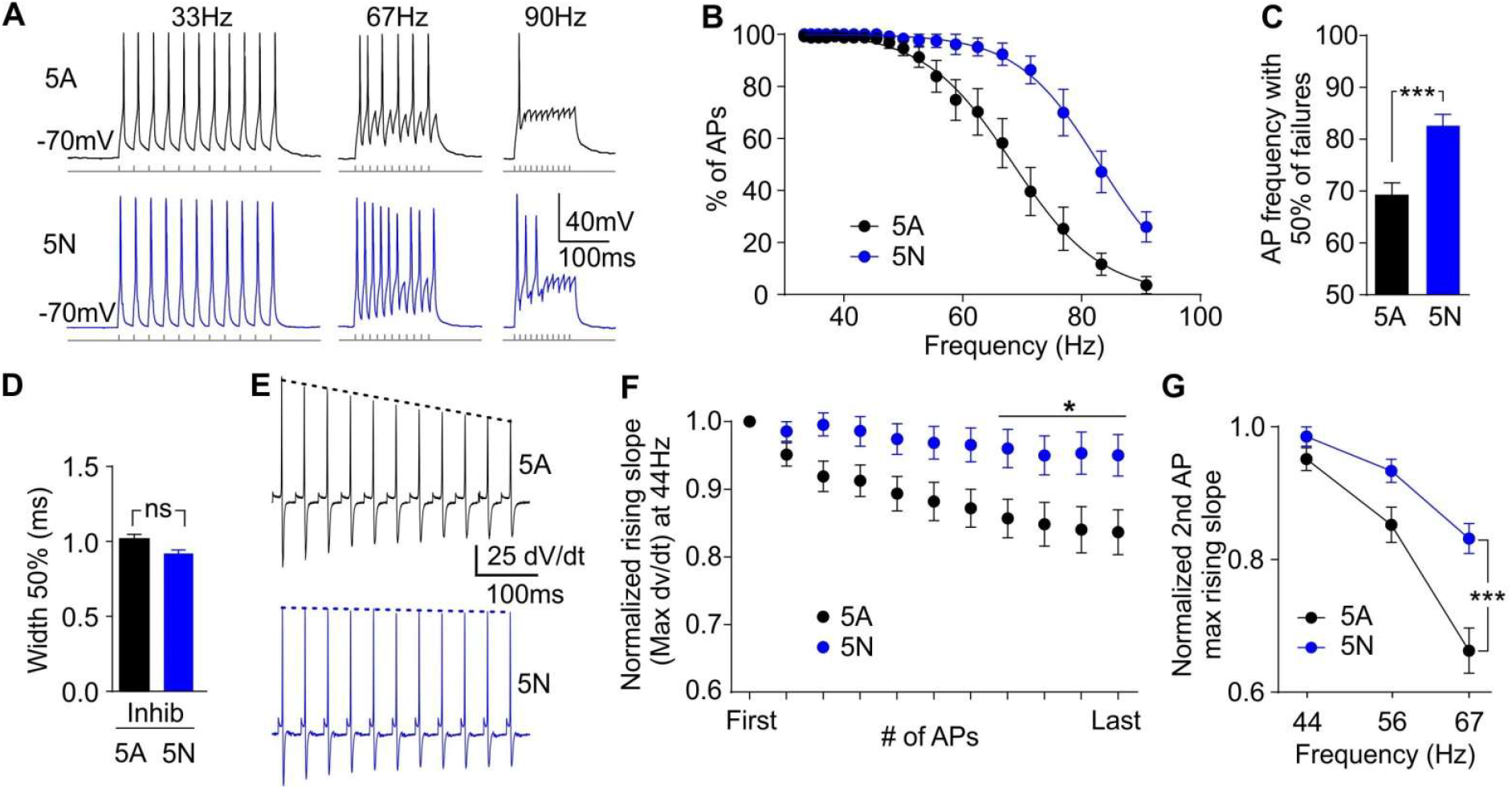
Splicing in Nav1.1 is sufficient to alter spike reliability of interneurons during rapid trains. **(A)** Representative traces from interneurons transfected with adult (top, black) and neonatal (bottom, blue) variants of Nav1.1 showing reduced ability of the A variant to support action potentials (APs) at higher frequencies. **(B)** AP failures increase with stimulation frequency. **(C)** The frequency at which interneurons fired APs at 50% of stimuli was significantly higher for neurons expressing N channel variants (N, n = 12; A, n = 12; *** P = 0.0005, unpaired two-tailed Student’s T test). **(D)** Splicing in Nav1.1 does not change the half width of APs in interneurons. The mean half width was similar for interneurons expressing N and A variants (ns P > 0.05, unpaired two-tailed Student’s T test). **(E)** Representative second differentials of the APs evoked by a series of steps from interneurons expressing A (top, black), and N (bottom, blue) splice variants showing slowing of the rising phase of adult only. The height of the peak, as indicated by the dotted lines, corresponds to the steepest part of the rising slope. Traces are from stimuli at 44 Hz, which is the fastest rate of stimuli where interneurons expressing A variants were able to support APs for all steps. **(F)** A splice variants show reduced ability compared to N variants to maintain fast rising slopes of APs during trains of stimuli. By the end of the series the APs supported by the A variants were significantly slower than those supported by N variants (* P < 0.05; two-way ANOVA followed by Bonferroni correction for multiple comparisons, A n = 12; N n = 13). **(G)** At higher rates of firing the reduction in rising slope for A variants is evident after a single AP. Cells which failed to fire APs were excluded (A n = 12; N n = 13). The difference was significant at 66 Hz (*** P < 0.001; Two-way ANOVA followed by Bonferroni correction for multiple comparisons).

Finally, inclusion of the A exon of Nav1.1 led to a substantial slowing in the rising phase of action potentials in inhibitory neurons exposed to repetitive stimuli (Figure 3E, F), which was greater than the effect of splicing of Nav1.2 in excitatory neurons (Figure 2E, F). While interneurons expressing the A variant of Nav1.1 had a pronounced slowing of the rising slope as trains of stimuli progressed (similar to either variant of Nav1.2 in excitatory neurons), interneurons expressing the N variant maintained fast rising slopes throughout the train of pulses. This preservation of the fast rising slope is consistent with sustained sodium channel availability when the N variant is expressed.

These data indicate that, although splicing of sodium channels has conserved effects in HEK cells (altering availability after inactivation), it results in very different consequences on APs in distinct neuronal populations in which the channels dominate.

### Neuronal background filters the effects of sodium channel splicing

To determine whether the effects of splicing were dependent on channel biophysics or on the cellular background, we asked whether the effects of splicing of Nav1.1 in interneurons carry over to excitatory neurons. Specifically, do splice variants of Nav1.1 expressed in excitatory neurons behave like Nav1.1 in interneurons, or more like Nav1.2 variants in excitatory neurons? As previously, neurons expressing TTX-resistant splice variants of Nav1.1 had similar passive membrane properties (Table 3).

**Table 3.** Membrane properties of excitatory neurons transfected with splice variants of NaV1.1 (mean ± s.e.m.)

|  | **5A (n=19)** | **5N (n=13)** |
| --- | --- | --- |
| **RMP (mV)** | -49.00 ± 2.66 | -55.00 ± 2.81 |
| **Input resistance (MΩ)** | 188.00 ± 21.74 | 174.00 ± 16.22 |
| **V threshold (mV)** | -22.87 ± 1.22 | -23.33 ± 1.11 |
| **Current threshold (pA)** | 308.00 ± 31.64 | 265.50 ± 24.13 |
| **Rising slope (V/s)** | 68.74 ± 6.76 | 62.99 ± 10.58 |
| **AP Amplitude (mV)** | 41.10 ± 2.53 | 38.81 ± 3.69 |
| **Repolarizing slope (V/s)** | -42.79 ± 2.079 | -36.50 ± 3.46 |

When driven with repetitive depolarizing pulses, excitatory neurons expressing N and A variants of Nav1.1 showed similar failure rates (Figure 4A-C). Expression in excitatory neurons thus prevented variants of Nav1.1 from imposing the sustained rise times in trains of stimuli, instead variants of Nav1.1 behaved more like variants of Nav1.2. In these experiments, neuronal background thus predominates in determining whether splice variants of sodium channels are sufficient to change maximal firing rates. However, neuronal background was not sufficient to completely override the channel type, because in excitatory neurons splicing in Nav1.1 was also not sufficient to change the half width of APs (Figure 4D). This suggests that the effect of splicing on half width is specific to Nav1.2, and not due to cellular background. We conclude that splice variants have conserved effects at molecular level, but that the cellular backgrounds predominate in determining how these effects translate to changes in neuronal activity.

**Figure 4:**
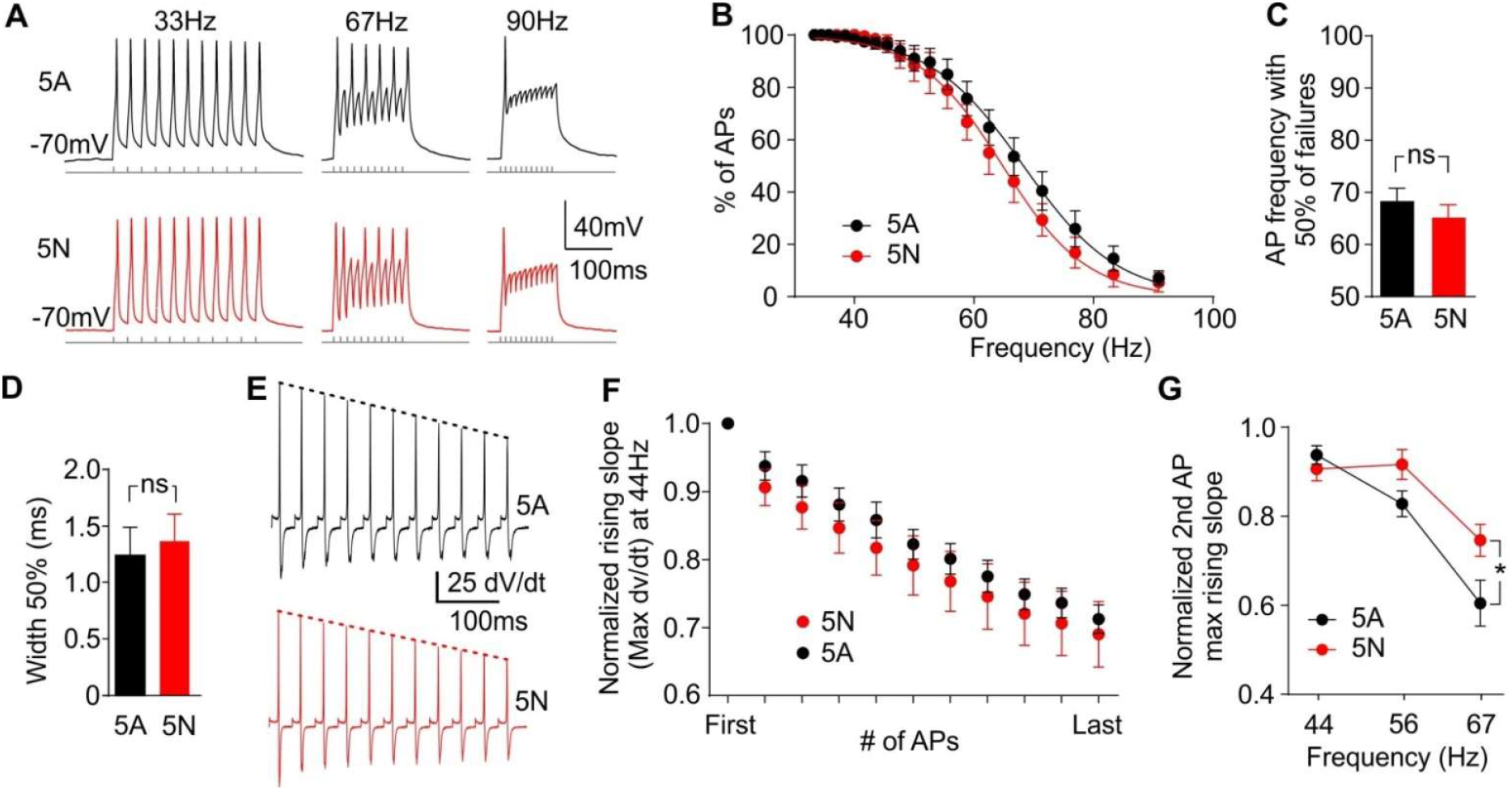
The effect of splicing is conserved, but consequences depend on cell background. **(A)** Representative traces from excitatory neurons transfected with A (top, black) and N (bottom, red) splice variants of Nav1.1. Recordings were carried out in 1μM TTX to block endogenous channels. **(B)** As with variants of Nav1.2, neither splice variant of Nav1.1 is able to sustain high rates of firing in these cells. **(C)** The rate at which neurons fired action potentials (APs) at 50% of stimuli, as calculated by the average of exponential fits to individual cells, were not different (ns P > 0.05, unpaired two-tailed Student’s T test). **(D)** Splicing in Nav1.1 does not change the half width of APs in excitatory neurons. The mean half width was similar for neurons expressing adult (1.54 ± 0.08 ms) and neonatal variants (1.36 ± 0.08 ms; ns P > 0.05, Mann-Whitney test). RMP, Input resistance, AP thresholds, maximal rising slope, amplitude, and maximal repolarizing slope were all similar for neurons expressing both variants. **(E)** The second differentials of APs evoked by a representative series of steps from an excitatory neuron expressing adult (top, black), and neonatal variants (bottom, red) both show decay in rate of rising slope. **(F)** During trains of stimuli both A and N variants of Nav1.1 show marked slowing of the rising phases of the APs. A and N splice variant rising slopes were similar throughout the trains. **(G)** At high firing frequencies the A variant of Nav1.1 showed reduced rising slope after a single AP. The difference is consistent with that seen in interneurons with Nav1.1 and excitatory neurons expressing splice variants of Nav1.2. Data show the change in rising slope as measured from the second derivative for excitatory neurons that fired in response to both the first and second stimuli. Cells which failed to fire APs were excluded (A, n = 10; N, n = 6). At 66 Hz the difference was significant (* P < 0.05; two-way ANOVA followed by Bonferroni’s multiple comparisons test).

### Delivery of epileptiform burst inputs highlights cell-type specific impact of splicing

Splicing in Nav1.1 has particular clinical relevance, as a polymorphism in this channel has been associated with response to AEDs and with the development of some forms of epilepsy (Kasperaviciute et al., 2013; Tate et al., 2005). This raises the possibility that splicing in Nav1.1 can modify how neurons respond during seizures, and the finding that the N variant sustains more action potentials without failures in trains of fast stimuli suggests these channels may also support greater activity during seizure activity. To test this possibility, we applied “activity clamp” (Morris et al., 2017), which is a dynamic clamp protocol that allows direct comparison of how different neurons respond to identical barrages of synaptic conductances recorded during an epileptiform event. It is especially informative of how individual neurons fire during seizures without contamination by the network consequences of neurotransmitter release from their terminals. Since Nav1.1 channels are thought to be particularly important in inhibitory neurons, we first asked whether splicing in Nav1.1 altered how interneurons responded during simulated seizure like inputs.

To reproduce the synaptic barrage, we recorded synaptic currents experienced by a representative neuron held in voltage clamp during epileptiform events (evoked by exposure to 4-Aminopyridine), and calculated the corresponding conductance waveforms (Figure 5, see methods). These were then delivered to neurons recorded in dynamic clamp with neurotransmitter receptors blocked and TTX applied (Morris et al., 2017). This method allowed us to probe how neurons transfected with different TTX-resistant sodium channel splice variants respond to synaptic inputs experienced during a seizure (Figure 5 - H).

**Figure 5:**
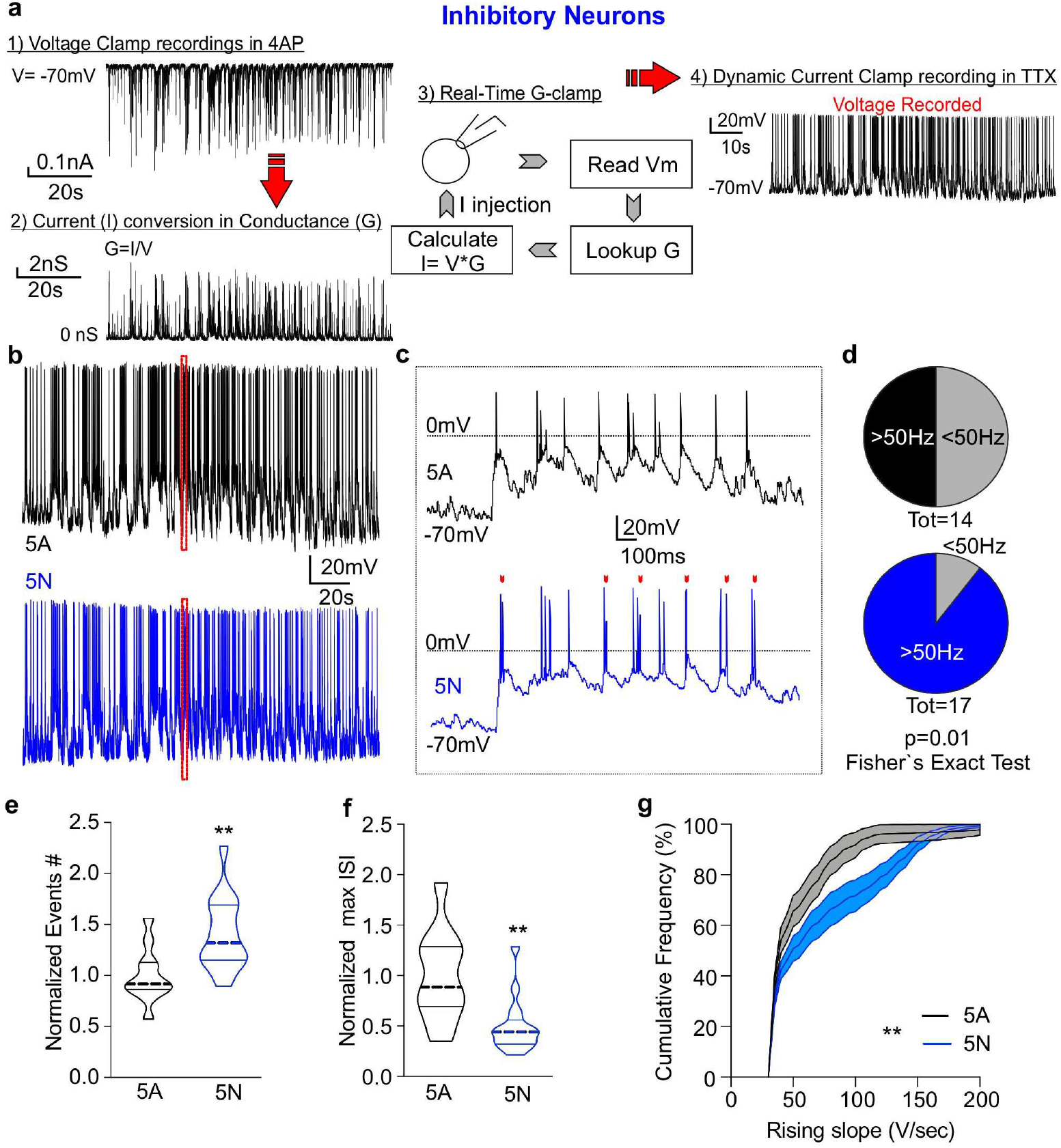
Dynamic clamp experiments demonstrate that splicing in Nav1.1 is sufficient to alter how interneurons respond during high frequency bursts. **(A)** An overview of the dynamic clamp procedure. Voltage clamp recordings of epileptiform bursts from interneurons in the presence of 4AP produced current traces (panel 1). To better capture cell type specificity of epileptiform activity, different inputs were used for inhibitory and excitatory recordings. Recorded current traces of synaptic inputs were converted via dynamic clamp to conductances (panel 2), which were used as a template for dynamic clamp (panel 3) to probe voltage responses (panel 4) from interneurons expressing the N or A splice variants of Nav1.1 in presence of TTX. **(B)** Representative voltage traces of the Nav1.1 variants resulting from dynamic current injection of the conductance trace shown in interneurons. **(C)** Enlarged representative traces of Nav1.1 variants from the dotted box in B. The red arrowheads indicate points where the N variant (blue) fired an AP and the A (black) failed. Note theses are generally during fast bursts. **(D)** Significantly more inhibitory neurons expressing the N variant attained firing frequencies above 50 Hz. **(E)** Number of AP (events) during activity clamp recordings normalized to each independent preparation (5A n=11; 5N n=16. **p<0.01 Unpaired Student’s t test). **(F)** Maximal inter-spike intervals reached during activity clamp recordings normalized to each independent preparation (5A n=11; 5N n=16. **p<0.01 Unpaired Student’s t test). **(G)** Cumulative plot representing the overall distribution of rising slopes during activity camp recordings (5A n=11; 5N n=16. **p<0.01. Splice variant factor, two-way ANOVA followed by Bonferroni multi-comparison).

Consistent with our current clamp data, interneurons expressing the A variant of Nav1.1 showed a lower maximal firing rate than interneurons expressing the N variant. Taking a cut-off for the instantaneous firing frequency of 50 Hz, 15/17 interneurons expressing the N variant exceeded this value, but only 7/14 interneurons expressing the A variant were able to fire at this frequency (Figure 5D; p = 0.019, Fisher’s exact test, two-tails). The number of APs, maximal frequency and rising slopes induced by activity clamp, were all significantly greater in interneurons expressing the N variant (Figure 5 E-G).

Our current clamp data suggest this effect may not be replicated, even using the same Nav1.1 splice variants, in excitatory neurons. To test this hypothesis, we expressed Nav1.1 variants in excitatory neurons, and delivered synaptic inputs recorded from an excitatory neuron during an epileptiform event (evoked by exposure to 4-Aminopyridine). As predicted, in excitatory neurons, neither the maximal spiking rate or the AP rising slope was not affected by expression of different splice variants of Nav1.1 (Figure 6 A-G). However, we detected a non-significant increase in rising slope (p=0.07), qualitatively similar to that observed with current clamp experiments in the same neuronal population (Figure 4 G and Figure 6 G). This difference between interneurons and excitatory neurons is consistent with the principle that the biophysics of sodium channel variants are particularly optimised for modulating activity in specific neuronal backgrounds, and Nav1.1 splicing is specifically tuned to modulate interneurons, but is ineffective in changing the maximal firing rate in excitatory neurons.

**Figure 6:**
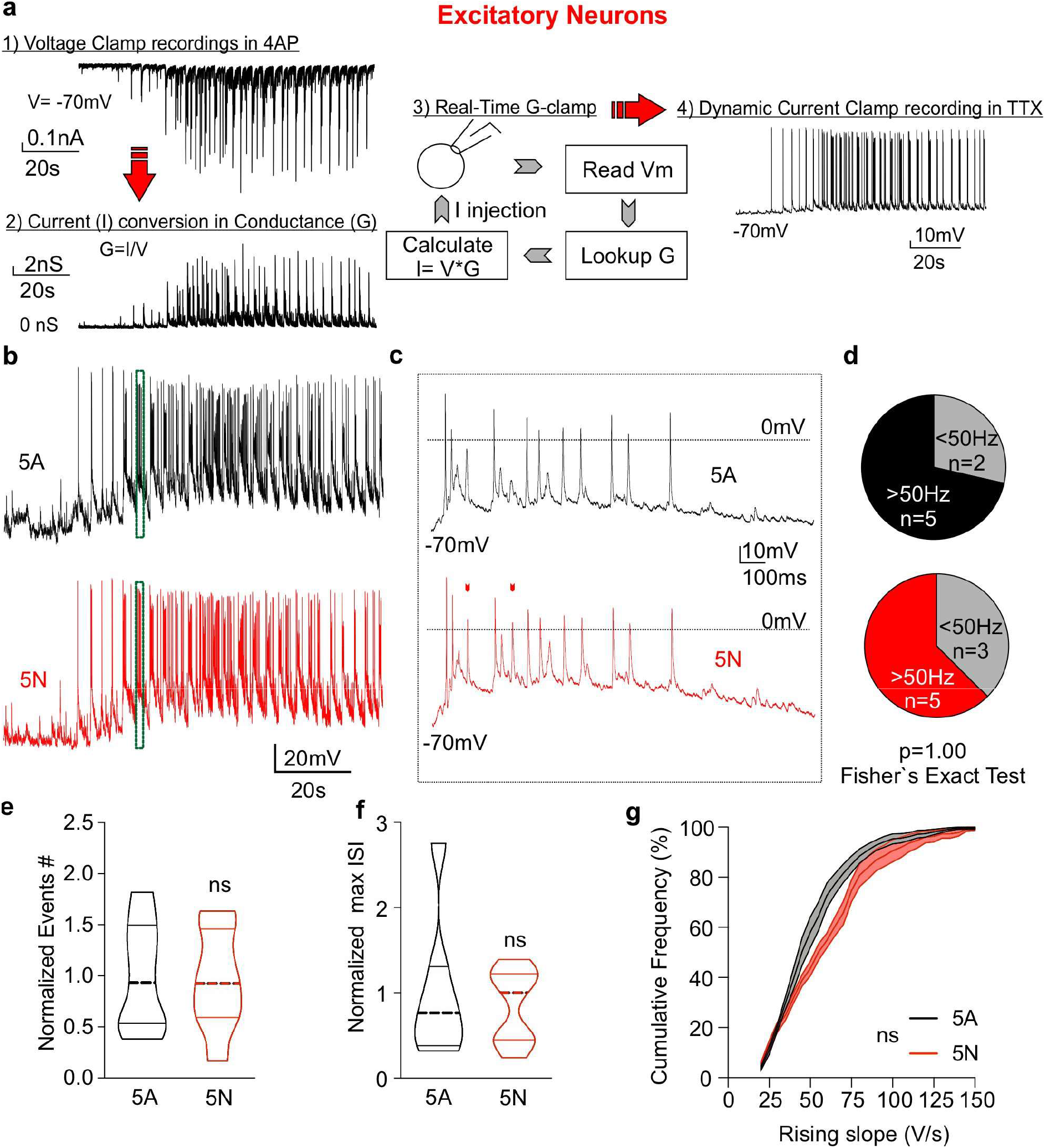
Dynamic clamp experiments demonstrate that splicing in Nav1.1 is not sufficient to alter how excitatory neurons respond during high frequency bursts. **(A)** An overview of the dynamic clamp procedure. Voltage clamp recordings of epileptiform bursts from excitatory neurons in the presence of 4AP produced current traces (panel 1). To better capture cell type specificity of epileptiform activity, different inputs were used for inhibitory and excitatory recordings. Recorded current traces of synaptic inputs were converted via dynamic clamp to conductances (panel 2), which were used as a template for dynamic clamp (panel 3) to probe voltage responses (panel 4) from excitatory neurons expressing the N or A splice variants of Nav1.1 in presence of TTX. **(B)** Representative voltage traces of the Nav1.1 variants resulting from dynamic current injection of the conductance trace shown in excitatory neurons. **(C)** Enlarged representative traces of Nav1.1 variants from the dotted box in B. The red arrowheads indicate points where the N variant (blue) fired an AP and the A (black) failed. Note theses are generally during fast bursts. **(D)** No difference in excitatory neurons expressing the N variant attained firing frequencies above 50 Hz. **(E)** Number of AP (events) during activity clamp recordings normalized to each independent preparation (5A n=7; 5N n=8. Unpaired Student’s t test). **(F)** Maximal inter-spike intervals reached during activity clamp recordings normalized to each independent preparation (5A n=7; 5N n=8. Unpaired Student’s t test). **(G)** Cumulative plot representing the overall distribution of rising slopes during activity camp recordings (P = 0.07; 5A n=11; 5N n=16. Splice variant factor, two-way ANOVA followed by Bonferroni multi-comparison).

The present data reveal that splicing in sodium channels is exquisitely tuned to the cell types in which these channels are found. Even the highly conserved consequences of splicing can have differing effects depending on the neuronal background. This specificity of splicing is seen in spite of the highly conserved site, and similar functional consequences when studied in non-neuronal cells.

## Discussion

The present study shows that the functional consequences of a conserved alternative splicing phenomenon are due both to the functional impact of the splicing per se on the sodium channel and to the cell type in which the channel subtype is expressed. The functional impact of splicing appears finely tuned to the type of neuron in which that channel dominates. These data show that sodium channels are so sensitive to neuronal environment, that even changes that produce similar biophysical effects can have different consequences for neuronal activity.

### The conserved effect of splicing on AP rise time and spike timing

Signal processing in the axon initial segment is highly dependent on the density of sodium currents (Kole and Stuart, 2012), consequently even by contributing relatively small changes to the availability of channels to pass currents, splicing could impact neuronal output. The main overall conserved effect of splicing is on the rise time of action potentials in trains. This could imply that splicing is important for modulating spike timing (Scott et al., 2014). It has also recently been shown that changing sodium channel availability, predominantly by sequestering channels into unstable inactivated states, allows channels to act as leaky integrators to control firing probability in neurons in response to previous activity (Navarro et al., 2020). The inactive states we have focused on are the fast inactive states, but our findings are consistent with the change in proportion of channels entering fast inactive states having effects on high frequency firing rates. In our conditions only variants of Nav1.1 in inhibitory neurons were sufficient to modify firing frequency, suggesting that the limits of frequency in excitatory neurons are set by different parameters, or other populations of channels, such as relatively few Kv3 channels in these cells (Gu et al., 2018).

### Splicing, channel availability in Nav1.1 and fast firing cells

In inhibitory neurons with small diameter axons, the density of sodium currents is even more critical, and is strongly correlated to speed of action potential propagation (Hu and Jonas, 2014). The proportionately greater effects of splicing in Nav1.1 on interneuronal firing are consistent with a subtle effect of this splicing on development of febrile seizures, and potentially dosage of AEDs (Tate et al., 2005). Unlike splicing in other sodium channel genes, splicing in SCN1A/Nav1.1 appears to be under evolutionary pressure to remove the N exon, and reduce the channels that allow the highest frequencies of firing (Liavas et al., 2017). The interaction between splicing and interneurons which fire rapid action potentials may be confounded by the presence of sodium channels that do not completely inactivate, but remain available, after these fast action potentials (Carter and Bean, 2011).

Although we have considered Nav1.1 as predominantly interneuronal, and Nav1.2 as predominantly expressed in (young) excitatory neurons, it should be noted this segregation is not complete. Indeed, recently Nav1.2 was found to play an important role in the AIS of a subset of interneurons (Li et al., 2014) highlighting how sodium channels can be specialized not just to different cell types, but to different regions within cells.

### Significance of splicing in Nav1.2

The effect on halfwidth of AP appears specialized for Nav1.2, and may reflect this channel’s importance during the development of neurons (Berecki et al., 2018; Gazina et al., 2015; Spratt et al., 2019). The roles of Nav1.2 are particularly important in back-propagating APs that invade the somatodendritic region (Spratt et al., 2019). Modulation of these currents by G-protein coupled receptors is sufficient to alter spike timing (Yu et al., 2018). Recently, reducing channel availability with either relatively low concentrations of TTX (20 nM) or phenytoin (100 μM) was shown to be critical for determining the non-linear amplification of excitatory inputs to dendrites (Hsu et al., 2018), suggesting that modulation of sodium channel availability, such as seen in splicing, may have robust consequences on signal integration in dendrites.

Splicing in Nav1.2 is closely linked to neuronal dysfunction in early development (Berecki et al., 2018; Thompson et al., 2020). In vivo, restricting Nav1.2 to only the A variant during development increases the excitability of layer 2/3 pyramidal neurons (Gazina et al., 2015). We did not observe a net difference in the firing properties of excitatory neurons in our experimental setting. This may be a reflection of the greater potential for homeostatic compensations during development in vivo, and the fact that in our system, in the absence of TTX, cells could express a range of sodium channels. Our data underline, in the absence of homeostatic changes, how sodium channel function has specific effects in different neuronal backgrounds, but do not give long term readouts of how cells might respond to being restricted to individual variants. However, in support of the neuronal parameters being sufficient to change the impact of splicing, in models based on excitatory neuronal parameters the A variants of Nav1.2 were sufficient to maintain a higher rate of firing (Thompson et al., 2020), albeit with a slightly different stimulation protocol than we used here. The observed differences in biophysical properties, and modelling are consistent with the exquisite sensitivity for sodium channels for the cellular background. Indeed, though splicing appears to have highly conserved effects on channels when they are expressed in the same type of cells, the consequences of these conserved effects are strongly divergent when translated to different neuronal environments.

Overall this work sets a precedent for the specificity of splicing in voltage gated channels and the precise tuning of these channels to cellular background.

## Acknowledgements

This work was funded by the Wellcome trust (WT104033AIA; 212285/Z/18/Z), the MRC (R/L01095X/1; MR/L003457/1). SS was funded in part by a Fellowship from the Royal Society (UF140596). GL was supported in part by a Marie Skłodowska-Curie Individual Fellowship, and by Epilepsy Research UK. AL was funded by an MRC PhD studentship.

